# The cyclic di-GMP network is a global regulator of phase-transition and attachment-dependent host colonization in *Erwinia amylovora*

**DOI:** 10.1101/2021.02.01.429191

**Authors:** Roshni R. Kharadi, Kayla Selbmann, George W. Sundin

## Abstract

Cyclic-di-GMP (c-di-GMP) is an essential bacterial second messenger that regulates the transition to biofilm formation in the phytopathogen *Erwinia amylovora*. The c-di-GMP system in *E. amylovora* is comprised of 12 diguanylate cyclase/Edc (dimerize cyclic-di-GMP) and phosphodiesterase/Pde (hydrolyze cyclic-di-GMP) proteins that are characterized by the presence of GGDEF and/or EAL motifs in their domain architecture. In order to study the global regulatory effect (without the inclusion of systemic regulatory impedance) of the c-di-GMP system in *E. amylovora*, we eliminated all 12 *edc* and *pde* genes in *E. amylovora* Ea1189Δ12. Comparisons between the representative transcriptomic profiles of Ea1189Δ12 and the combinatorial *edc* gene knockout mutant (Ea1189Δ5) revealed marked overall distinctions in expression levels for targets in a wide range of regulatory categories, including metabolic pathways involved in the utilization of methionine, isoleucine, histidine, etc. as well as critical signal transduction pathways including the Rcs phosphorelay and PhoPQ system. A complete loss of the cyclic-di-GMP signaling components resulted in the inability of Ea1189Δ12 cells to attach to and form biofilms in vitro and within the xylem vasculature in apple shoots. Using a flow-based in vitro biofilm system, we found that initial surface sensing was primarily dependent on the flagellar filament (FliC), following which the type IV pilus (HofC) was required to anchor cells to the surface to initialize biofilm development. A transcriptomic analysis of WT *E. amylovora* Ea1189 and Ea1189Δ12 cells in various stages of biofilm development revealed that cyclic-di-GMP based regulation had widespread effects on purine and pyrimidine biosynthesis pathways, amylovoran biosynthesis genes and the EnvZ/OmpR signal transduction system. Additionally, complementing individual eliminated genes back into Ea1189Δ12, and the collective evaluation of several virulence factors, enabled the correlative clustering of the functional effect rendered by each Edc and Pde enzyme in the system.

**Significance:** Cyclic-di-GMP dependent regulation, in the context of biofilm formation, has been studied in several bacterial systems. However, the comprehensiveness of the studies exploring the role of individual genetic components related to cyclic-di-GMP is affected by the often large number of diguanylate cyclase and phosphodiesterase enzymes present within individual bacterial systems. To explore the evolutionary dependencies related to cyclic-di-GMP in *E. amylovora*, we used a collective elimination approach, whereby all of the enzymes involved in cyclic-di-GMP metabolism were eliminated from the system. This approach enabled us to highlight the critical importance of cyclic-di-GMP in plant xylem colonization due to its effect on surface attachment. Additionally, we highlight the global transcriptomic effect of cyclic-di-GMP dependent signaling at various stages of biofilm development. Our approach is aimed at exploring the regulatory role of individual cyclic-di-GMP related enzymes in a background that is free from any redundancy-based feedback.

## Introduction

Bacterial cyclic di-GMP (c-di-GMP) networks consist of diguanylate cyclase (Dgc) enzymes that synthesize and phosphodiesterase (Pde) enzymes that hydrolyze bis (3’,5’)-cyclic diguanosine monophosphate (c-di-GMP), and downstream c-di-GMP receptors, usually either proteins or riboswitches that bind c-di-GMP and transduce the signal into either a regulatory response or a functional response in the case of allosteric activation upon c-di-GMP binding. The number of enzymes that are involved in c-di-GMP turnover in bacteria from the order Enterobacterales can be as many as 34 or as few as (1). Cyclic di-GMP networks in bacteria are most commonly known to regulate transitions from a motile to a sessile state or from an acute infection state to a more chronic, resilient state (2). Since many Dgc and Pde enzymes have external domains that are responsive to environmental signals, the c-di-GMP network represents a critical system through which bacteria can utilize an environmental assessment to drive significant alterations in lifestyle.

The dimerization of c-di-GMP from GTP substrates is mediated by Dgc enzymes, often marked by the presence of a GGDEF domain (3). The hydrolysis of c-di-GMP, which yields either the linear nucleotide pGpG (5’-phosphoguanylyl-(3’→5’)-guanosine) can be mediated by c-di-GMP-specific Pde enzymes that are characterized by the presence of an EAL motif or a HD-GYP motif (4-6). Finally, the HD-GYP class of Pdes and oligoribonucleases can further hydrolyze pGpG into GMP subunits (6, 7). In *E. amylovora*, five active Dgcs (EdcA-E) and three active Pdes (PdeA-C) have been functionally characterized (8, 9). As documented in other bacterial systems including *Escherichia coli* (10), *Pseudomonas aeruginosa* (11), *Salmonella enterica* (12), and *Vibrio cholerae* (13), the presence of a large number of Dgc and Pde enzymes is a common occurrence, as functional redundancy is often observed during analyses of these enzymes.

The primary approach used to study the regulatory effect of individual components of c-di-GMP turnover is by deleting one or more genes in combination and evaluating phenotypic and regulatory changes. Among these studies, notable ones that have been aimed at characterizing the roles of a large number of Dgc and/or Pde enzymes include Solano et al. (12) wherein a multigene mutant that eliminated 12 *dgc* genes in *Salmonella* was generated. Restoring critical residues in this mutant revealed that STM4551 was a regulator of multiple virulence factors independent of its enzymatic activity towards c-di-GMP production, and four other Dgcs were found to enzymatically regulate cellulose synthesis in a c-di-GMP dependent manner (12). Another study by Sarenko et al. (10) involved the systematic deletion of each one of the 29 individual Dgc and Pde encoding genes in *E. coli* and studying their effect on biofilm formation as well as EPS production. Similar to Solano et al. (10), this study also found that the functional effect of specific Dgcs and Pdes on virulence phenotypes could be either dependent or independent of their enzymatic effect on modulating c-di-GMP levels. Hybridization assessment revealed the presence of several subsets of Dgc and Pde components that were interconnected and shared regulatory interactions. Additionally, certain dominant Pdes (PdeH) were capable of counteracting the effect of multiple Dgcs, by affecting the intracellular levels of c-di-GMP (10).

While the targeted elimination of one or more genes in the c-di-GMP regulatory system is an easy approach that can highlight significant changes in the regulation of critical virulence factors, the presence of multiple other genes in a highly redundant system can mask some of the effects occurring due to the loss of one particular gene, unless their enzymatic activity is particularly high compared to other similar components. Additionally, although the aforementioned studies have been able to explore the c-di-GMP dependent and independent effect of specific Dgcs and Pdes, the overall regulatory effect of each of these enzyme classes has not been explored in a background that does not include any additional impedance/interaction with any of the other components in the network. The retention of such a high number of Dgcs and Pdes in bacterial pathogen systems raises questions about the evolutionary significance of developing multilayered genetic control strategies.

In order to address these concerns and with the overarching aim of exploring the role of the global effect of c-di-GMP on virulence manifestation in the host, we identified and eliminated all genes that encoded for proteins with a GGDEF and/or EAL motif in the plant pathogen *Erwinia amylovora*, the causal agent of fire blight disease of rosaceous plants (14). An evolutionary adaptation that helps *E. amylovora* systemically colonize the apple host is its ability to attach to the walls of xylem vasculature and form robust biofilms within the xylem channels, thus enabling extensive proliferation of the pathogen during this stage of the disease cycle (15). Cyclic-di-GMP is one of the critical factors that regulates biofilm formation in *E. amylovora* (8, 9), and elevated intracellular levels of c-di-GMP have been correlated with increased levels of biofilm development in static and flow based in vitro systems (9, 16). The exopolysaccharides amylovoran and cellulose are the primary extracellular matrix components that lend structural integrity to *E. amylovora* biofilms within the host (15, 17). Amylovoran synthesis and secretion is dependent on the 12-gene *ams* operon, with *amsG* being the first gene in the operon (15). *amsG* transcription is positively regulated by c-di-GMP in *E. amylovora*, and this has an overall impact on amylovoran production in vitro and in bacterial ooze production in an immature pear fruit infection model (9). Cellulose production is activated by c-di-GMP through allosteric binding to BcsA (the catalytic subunit of cellulose) (17). In addition to biofilm formation, c-di-GMP also negatively regulates type III secretion (T3SS) mediated virulence via the transcriptional downregulation of *hrpL* (alternate sigma factor required for the transcription of *hrp* genes) and a reduction in the translocated levels of the T3SS effector, DspE (pathogenicity factor) (8, 9).

In this study, we eliminated 12 genes of the c-di-GMP network in *E. amylovora* Ea1189 (Ea1189Δ12), including five Dgcs, four Pdes, and three proteins that contain degenerate GGDEF and/or EAL domains, resulting in an *E. amylovora* strain with no background c-di-GMP formative, degradative, or signaling activity. Additionally, we used the complete diguanylate cyclase gene deletion mutant Ea1189Δ5 to compare and contrast phenotypic and transcriptomic profiles relative to Ea1189Δ12. We also explored the ability of *E. amylovora* Ea1189Δ12 to colonize its host and evaluated general surface interaction mechano-dynamics in the context of biofilm development. Further, we used a transcriptomic approach to explore the global regulatory effects of c-di-GMP during various stages of biofilm initiation and development, and conducted an aggregative evaluation of the effect of each of the twelve enzymes on an array of virulence factors in *E. amylovora*.

## Results

### *E. amylovora* has an array of twelve proteins with GGDEF and/or EAL motifs

The *E. amylovora* genome (ATCC-49946) contains 12 genes that encode for proteins that have a GGDEF and/or EAL motif (as annotated by Pfam v 33.1) (18, 19). Proteins marked by HD-GYP motifs are absent in *E. amylovora* (9). The list of 12 proteins along with their domain architecture is graphically mapped in Figure 1. EdcA-E have been previously mapped and characterized by Edmunds et al. (8). The functional characteristics regulated by PdeA-E have been studied by Kharadi et al. (9). EAM_3378, EAM_3136 (CsrD), EAM_2449 and EAM_1579 have been primarily documented in this study. The N-terminal end of all proteins except EdcA, EAM_3378 and EAM_1579 include a wide array of periplasmic sensory domains and multiple transmembrane helices (as predicted by TMHMM Server v 2.0) (20). EdcA, PdeC and CsrD contained both GGDEF and EAL domains (Figure 1).

**Figure 1:**
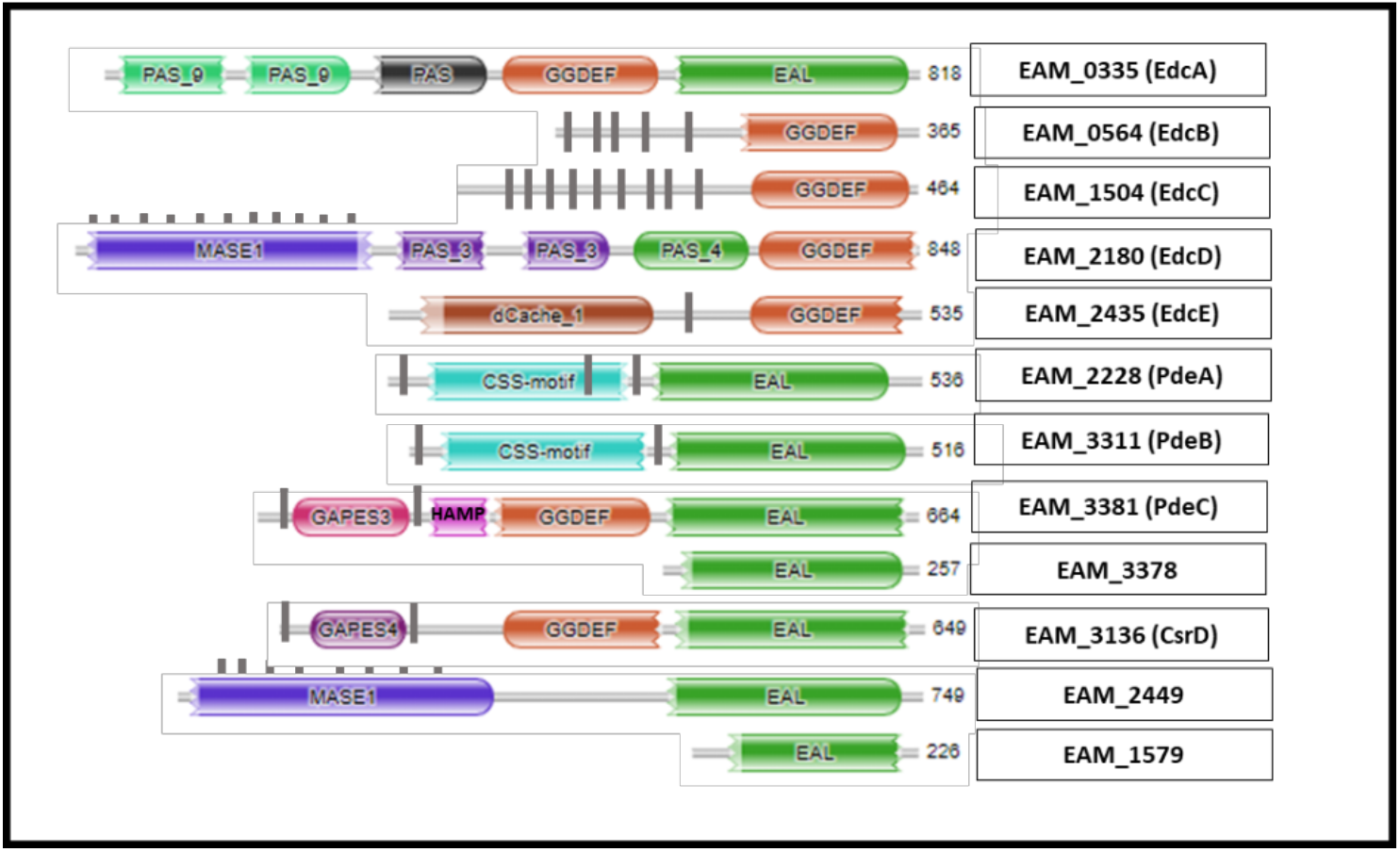
A graphical representation of the domain architecture of all twelve proteins with predicted GGDEF and/or EAL domains present in the *E. amylovora* genome (Accession ATCC-49946) (18). No proteins containing HD-GYP domains were bioinformatically predicted to be present in the genome. Pfam (v. 33.1) (19) was used to map the domain arrangement for each of the residues. Filled gray bars along the length of the protein represent Transmembrane domains as predicted by TMHMM Server (v. 2.0) (20). The proteins have been identified here using the annotated gene IDs (Accession ATCC_49946). Additionally, the previously published modified annotations for a subset of the proteins are mentioned in parenthesis adjacent to the gene IDs.

### Ea1189Δ12 and Ea1189Δ5 are characterized by marked distinctions in their overall transcriptomic profiles

A comparative transcriptomic profile of Ea1189Δ5 compared to Ea1189Δ5 revealed a total of 398 differentially expressed genes (DEGs) (Figure S1A). The greatest gene ontology (GO) based pathway enrichment for both the up- and down-regulated DEGs comprised of metabolic pathways including organic substrate utilization for histidine, isoleucine and methionine. Other significantly represented pathways comprised of organic substrate and peptidoglycan biosynthesis, and active transport pathways including putrescine, arabinose and ferrichrome ABC transport genes (Figure S1B). Confirmatory gene expression values for representative genes is presented in Figure S2A.

### Surface attachment is dependent on c-di-GMP in *E. amylovora*

An initial assessment of biofilm formation under flow in *E. amylovora* (using GFP labelled cells) revealed a marked absence in the detection of any fluorescence signal indicating attachment or biofilm development in Ea1189Δ12 compared to WT Ea1189 (Figure S3). However, restoring the individual genes *edcA-E* which contribute to c-di-GMP production (Figure S3) led to the restoration of biofilm development, although relatively sparse compared to WT Ea1189. In order to confirm that the source of the impediment to biofilm development in Ea1189Δ12 was the lack of initial surface attachment, we monitored the interaction of cells with the base of the flow chamber upon initial contact. Figure S4 (A and B) presents collated images in the form of a time lapse video presenting the dynamics of surface interaction of Ea1189 and Ea1189Δ12 cells introduced into the chamber. Several Ea1189 cells were found to approach the basal portion of the flow cell and a subset of them would get attached in every frame of the video (Figure S4A). Over time, this led to a saturation of the image frame with even GFP signals from attached cells. In contrast, Ea1189Δ12 cells were found to recurringly approach the basal surface, but failed to attach to surface irreversibly. Due to the pace of the video, this occurrence plays out in the form of momentary GFP signal increases and lapses as cells approach and then reproach from the surface (Figure S4B). Following the 1h allowed attachment period for both strains, the flow chambers were flushed and imaged using a z-stack series. Ea1189 cells retained widespread attachment, amplified in signal due to the merger of the z-stacked images. Ea1189Δ12 cells were unable to attach to the flow chamber surface (Figure 2).

**Figure 2:**
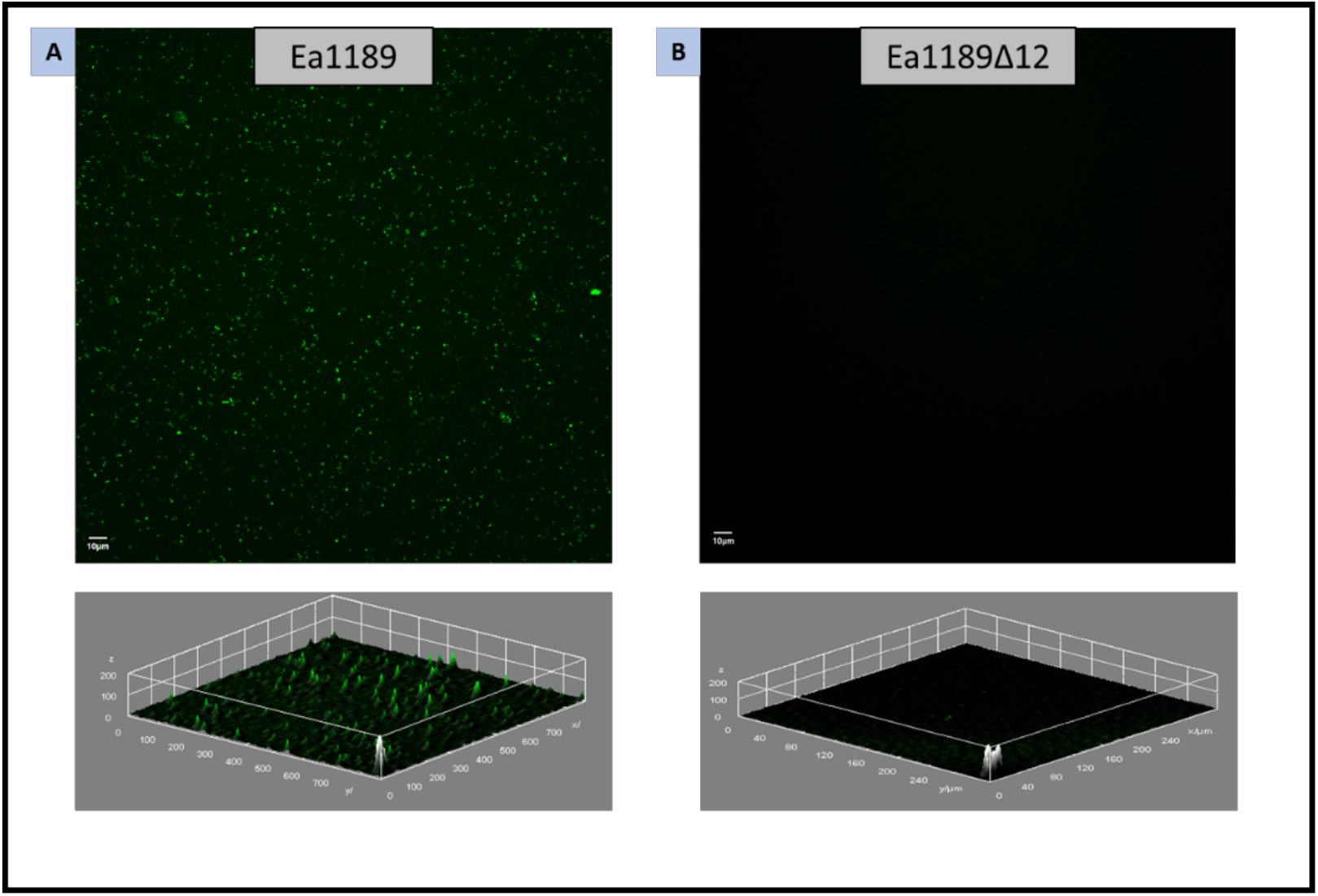
Z-stacked confocal microscopy images and concurrent signal intensity profiles showing the overall attachment occurring within the flow chamber 1 hr after the introduction of either *E. amylovora* Ea1189 (A) or Ea1189Δ12 (B) cells into the chamber, followed by the flushing of the chamber with 0.5X PBS. Ea1189 cells displayed widespread even attachment with interspersed patches of elevated fluorescence signal indicating potential multilayered attachment. Ea1189Δ12 cells failed to attach to the chamber surface.

### Type IV pilus regulates surface attachment in a c-di-GMP dependent manner in *E. amylovora*

In order to investigate which physical attachment appendage potentially regulates c-di-GMP mediated surface attachment, we utilized the hyper-attaching strain Ea1189Δ*hfq* (21). The flagellum, type IV pilus, fimbriae and curli fimbriae were targeted for mutations through the deletion of *fliC, hofC, fimA* and *crl* genes respectively (22). The Δ*hfq*Δ*fliC* and Δ*hfq*Δ*hofC* double mutants showed the greatest reduction in attachment relative to Ea1189Δ*hfq* (Figure S5). The overexpression of *hofC* but not *fliC* in Ea1189Δ12 restored attachment, however, the overexpression of *hofC* in Ea1189Δ12Δ*fliC* did not lead to any increase in attachment relative to Ea1189Δ12 (Figure 3). Additionally, the overexpression of *fliC* in Ea1189Δ12 did not yield any changes in the level of surface attachment (Figure S5).

**Figure 3:**
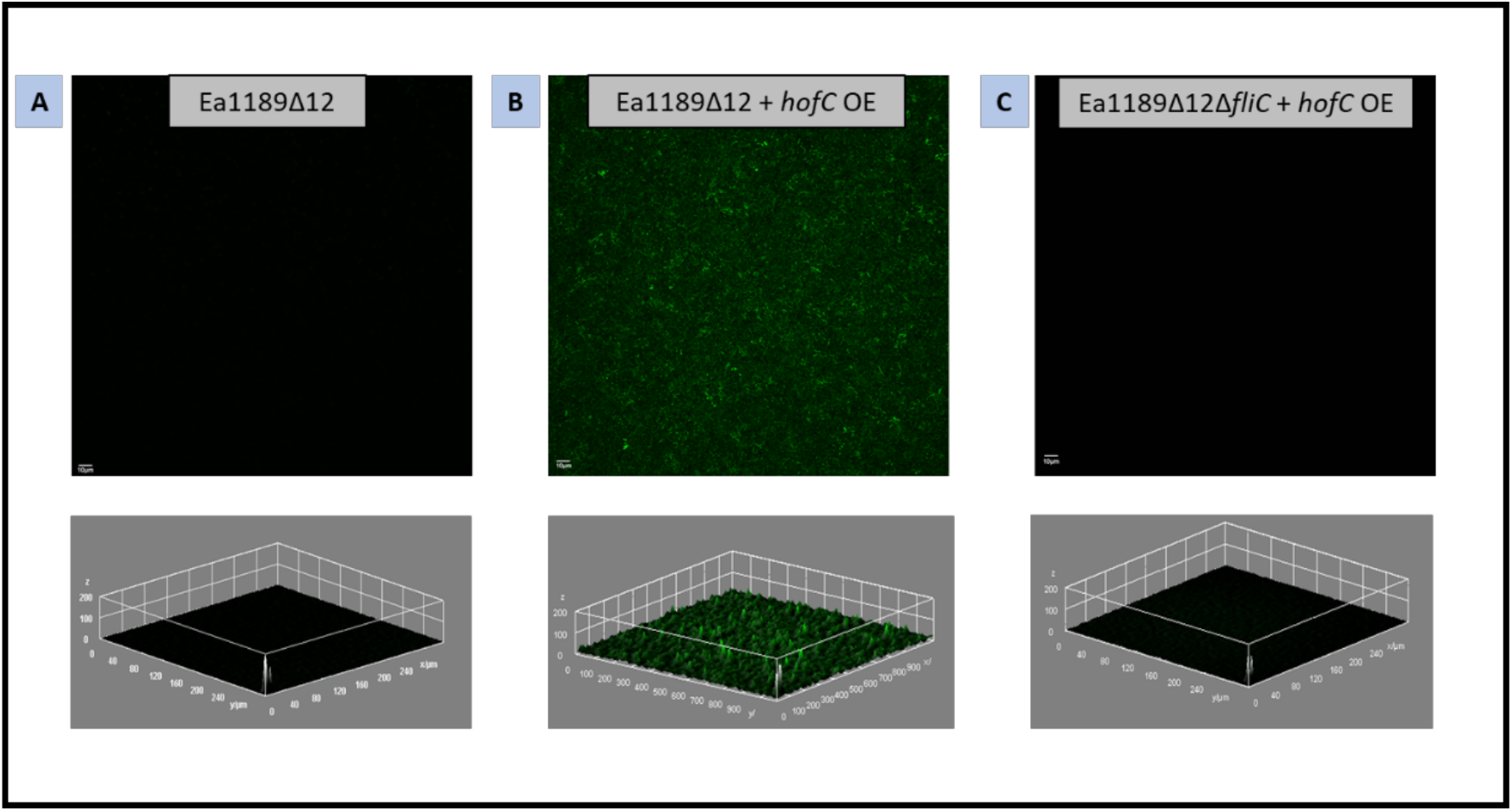
Z-stacked confocal microscopy images and concurrent signal intensity profiles showing the overall attachment occurring within the flow chamber one hour after the introduction of GFP labelled *E. amylovora* Ea1189Δ12 (A) and Ea1189Δ12 cells overexpressing *hofC* in the presence (B) and absence of *fliC* (C) followed by the flushing of the chamber with 0.5X PBS. While the over expression of *hofC* resulted in the occurrence of attachment within the chamber, this effect was negated upon the deletion of *fliC*.

### Xylem colonization by *E. amylovora* in the apple host is dependent on c-di-GMP

To check if the inability of Ea1189Δ12 cells to attach to a surface in vitro would translate to the attachment within vascular tissue, young apple leaves inoculated with Ea1189 and Ea1189Δ12 were evaluated using SEM to check for relative levels of proliferation and colonization in the apoplast and the xylem vessel channels. Ea1189 cells were detected abundantly in the apoplast 5dpi. In the xylem vessel channels, Ea1189 cells were able to attach and develop extensive biofilms and form an abundance of EPS, thus, functionally impeding the channels (Figure 4). In contrast, the apoplast region of leaves infected with Ea1189Δ12 showed relatively few pockets of cells with some extracellular material, whereas the xylem vessels showed no signs of colonization and no cellular attachment, EPS generation or biofilm formation was detected (Figure 4).

**Figure 4:**
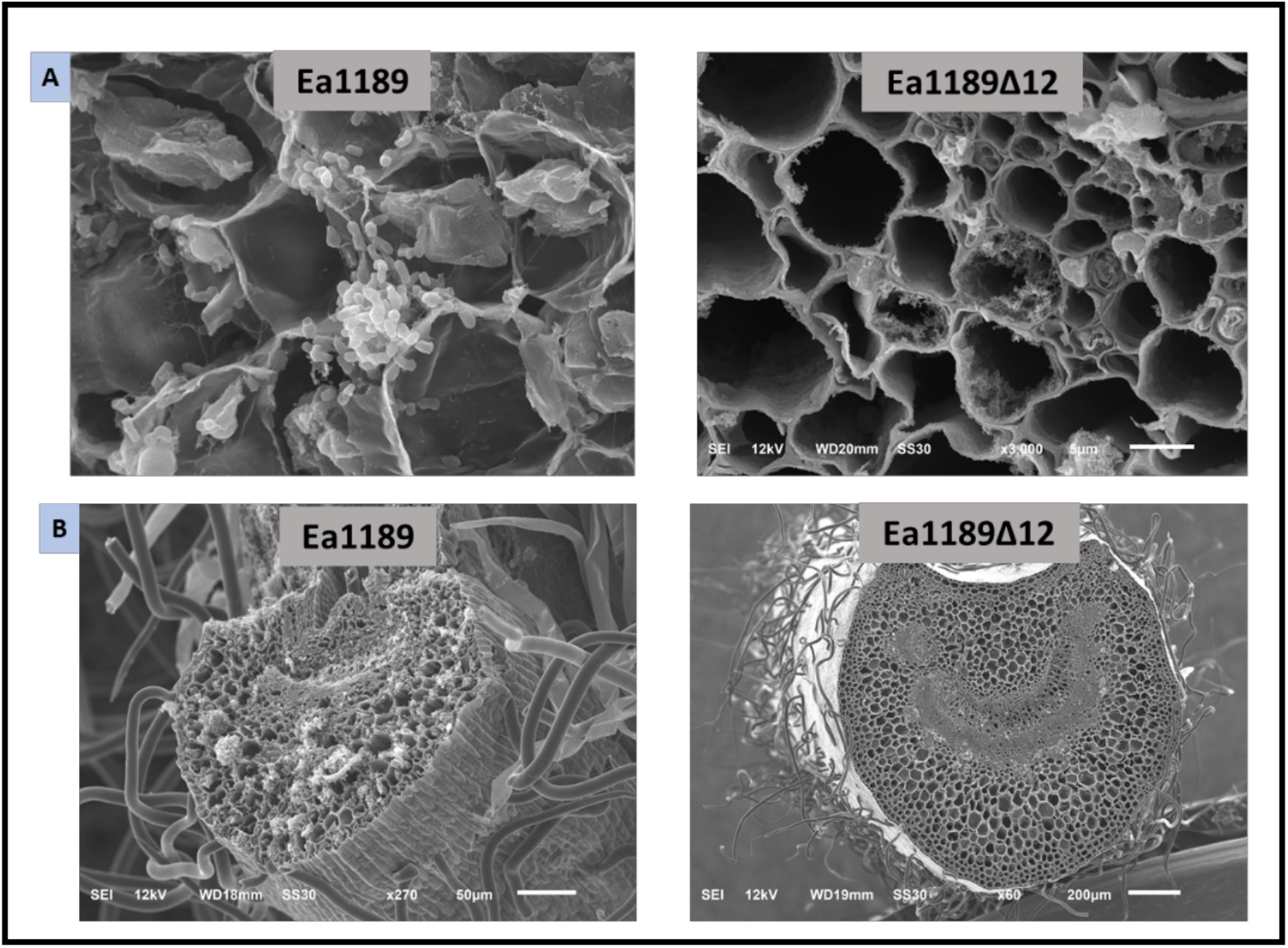
SEM images displaying the extent of colonization by *E. amylovora* Ea1189 and Ea1189Δ12 cells within apple leaf apoplasts (A) and petioles (B) 5dpi. WT Ea1189 was found to be able to proliferate within the apoplast and form extensive biofilms in the xylem channels of the petiole. Ea1189Δ12 shows very weak colonization in the apoplast (mainly marked by the presence of some extracellular materials) and no attachment or biofilm formation within xylem vessels.

### C-di-GMP transcriptionally regulates several critical pathways during different stages of biofilm development

An overall comparison of the differentially expressed genes (DEGs) among the cross-comparison of sample pairs revealed that cells exposed to the surfaces in a flow cell chamber and collected for RNA-Seq analysis (referred to as the ‘Unbound’ samples) showed the highest relative transcriptomic change (876 DEGs) for Ea1189Δ12 compared to Ea1189 (Figure 5A). For the comparison metric Ea1189Δ12 vs. Ea1189, the most representative functional enrichment of up and down regulated gene sets included several metabolic pathways, including sulfur, purine, carbon, cystine and pyrimidine metabolism. Additionally, several critical two component systems including the ArcAB, RcsABC, Bar/Uvr, PhoPQ and OmpR/EnvZ were among the down-regulated genes categories. (Figure 5C). A distribution tree represents the specific gene enrichment pathways for this comparison metric in Figure 5C. Figure S6 represents the categorical clustering of DEGs across the four sample types (Ea1189_Unbound, Ea1189Δ12_Unbound, Ea1189_Bound and Ea1189_Biofilm). A few of the notable categories of pathway enrichment across the four representative clusters of DEGs include diverse metabolic pathways including glutamine, arginine and isoleucine synthesis, flagellar biosynthesis, two-component systems including the Rcs phosphorelay system and the EnvZ/OmpR system, homologous recombination and the TCA cycle (Figure S6). Overall, significant transcriptomic shifts were recorded in the absence of c-di-GMP (Ea1189Δ12_Unbound) compared to conditions where c-di-GMP was present (Ea1189_Unbound, Ea1189_Bound and Ea1189_Biofilm). Additionally, in Ea1189, transcriptional patterns were contrastingly different within the three conditions tested, with the Ea1189_Unbound transcriptomic profile (including only DEGs) showed a relatively distinct pattern compared to the Ea1189_Bound and Ea1189_Biofilm stages (Figure S6). Representative genes targets were analyzed and compared to the transcriptomic data to validate the study (Figure S2).

**Figure 5:**
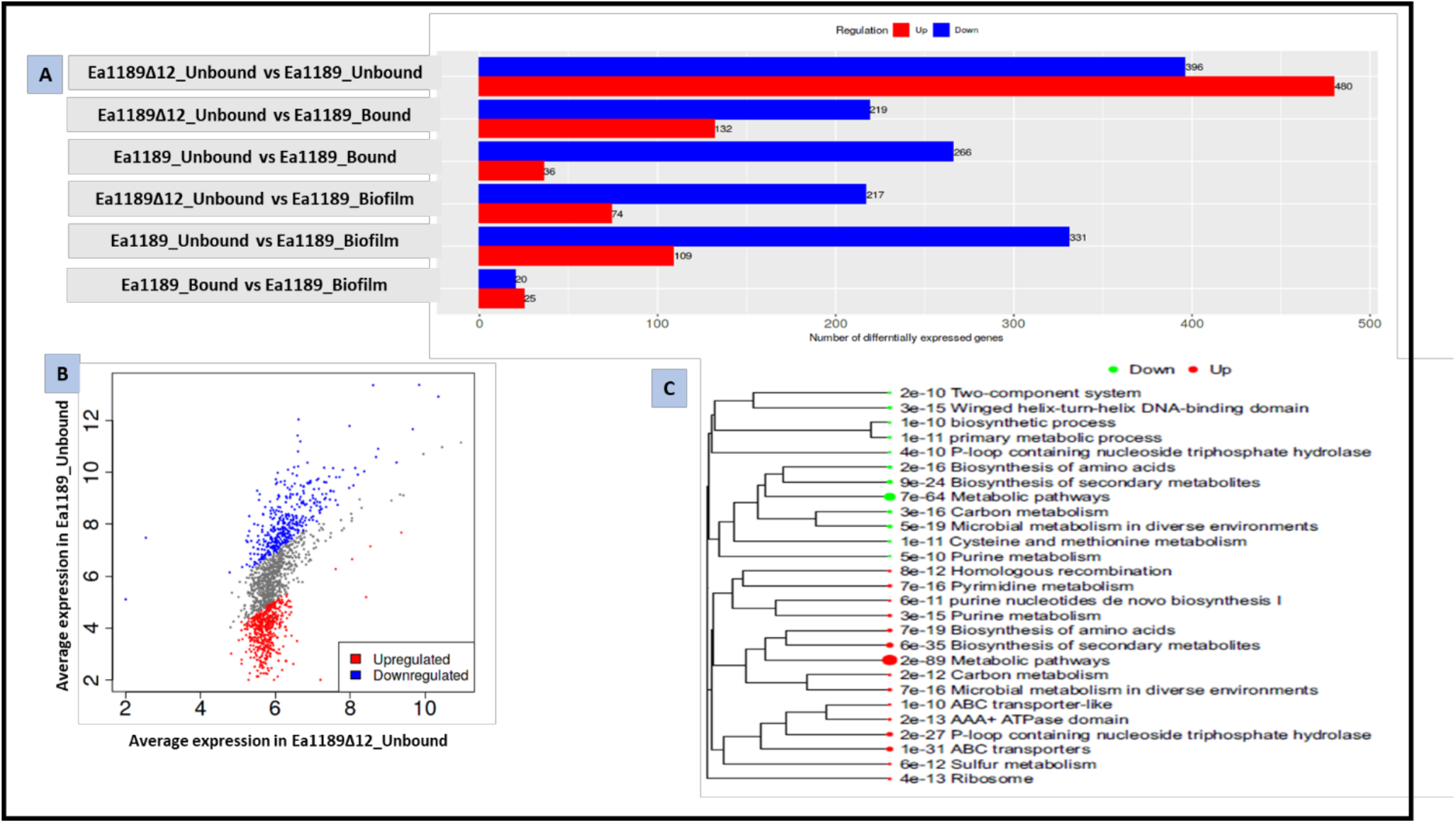
A) Numerical comparisons of the upregulated and downregulated DEGs evaluated via RNA-Seq across different comparative combinations comprising of four primary treatment/strain variants, *E. amylovora* Ea1189_Unbound, Ea1189_Bound, Ea1189_Biofilm and Ea1189Δ12_Unbound. B) A scatter plot displaying the expression trends of upregulated and downregulated genes when Ea1189_Unbound and Ea1189Δ12_Unbound were compared. C) A distribution tree displaying the gene ontology enrichment detected among upregulated and downregulated genes for Ea1189_Unbound vs. Ea1189Δ12_Unbound datasets. The proximity of the sub-branches of the tree indicates gene set similarities and the relative size of the filled circle represents the numerical proportion of DEGs in each pathway.

### Individual Dgc and Pde enzymes show differential regulatory patterns for key virulence determinants

An aggregate data analysis of key virulence factors in Ea1189, Ea1189Δ5, Ea1189Δ12 and Ea1189Δ12 restored with each individual deleted genes showed variable regulatory patters across the different parameters that were assessed (Figure 6). Evaluated post clustering (Pearson correlation), as expected, the regulatory patterns for the biofilm formation data clustered with *amsG* expression, amylovoran production, intracellular c-di-GMP levels and virulence in the apple shoot model. *hrpL* expression was found to cluster with the virulence levels recorded in the pear infection model (Figure 6A). Ea1189Δ5 clustered as the farthest relative outgroup within the matrix, compared to WT Ea1189, Ea1189Δ12 and the single gene restorative mutants. Ea1189Δ5 showed a marked contrast from Ea1189Δ12 in phenotypic patterns with a consistent pattern of downregulation in all evaluated parameters, including *hrpL* expression and virulence in the pear infection model, which was particularly distinct from Ea1189Δ12 (Figure 6A and 6B). Ea1189Δ12 restored with *edcA*-*E* showed increased correlation as compared to complemented mutants for rest of the genes. A strong factor impacting this observation is that EdcA-E are the only enzymes that were found to contribute to the physical generation of c-di-GMP. As a corollary, the data patterns for amylovoran production, *amsG* expression and biofilm formation indicated that the restoration of each of the *edc* genes had a positive impact on these characteristics. In terms of virulence regulation, Ea1189Δ12 showed elevated *hrpL* transcript levels and among the *edc* genes, the restoration of *edcE* led to the most decline in this metric (Figure 6A). The virulence patterns differed in the immature pear and apple shoot systems for the evaluated strains. The tissue necrosis levels in immature pear fruit closely resembled the patterns of *hrpL* transcription, whereas, in the apple shoot model, among the *edc* genes, only the restoration of *edcE* was able to cause a significant increase in virulence levels. Ea1189Δ12+*csrD* showed interesting regulatory patterns across the different parameters. While this gene did not contribute to c-di-GMP production, it had a positive impact on *amsG* transcription, biofilm formation and in terms of overall virulence levels (Figure 6A). A correlation matrix across the evaluated strains revealed a general cluster bifurcation between Ea1189Δ12 + *edcA-E* and the rest of the compliments. Interestingly, despite the previously documented functional redundancy and phenotypic homology observed among *pdeA-C*, the correlation based clustering was largely mixed (Figure 6B). Finally, our experimental approach also revealed new virulence based regulatory patterns for the previously uncharacterized genes EAM_3378, *csrD*, EAM_2449 and EAM_1579. Figures S7-S13 present the data analyzed for overall correlational matrix analyses.

**Figure 6:**
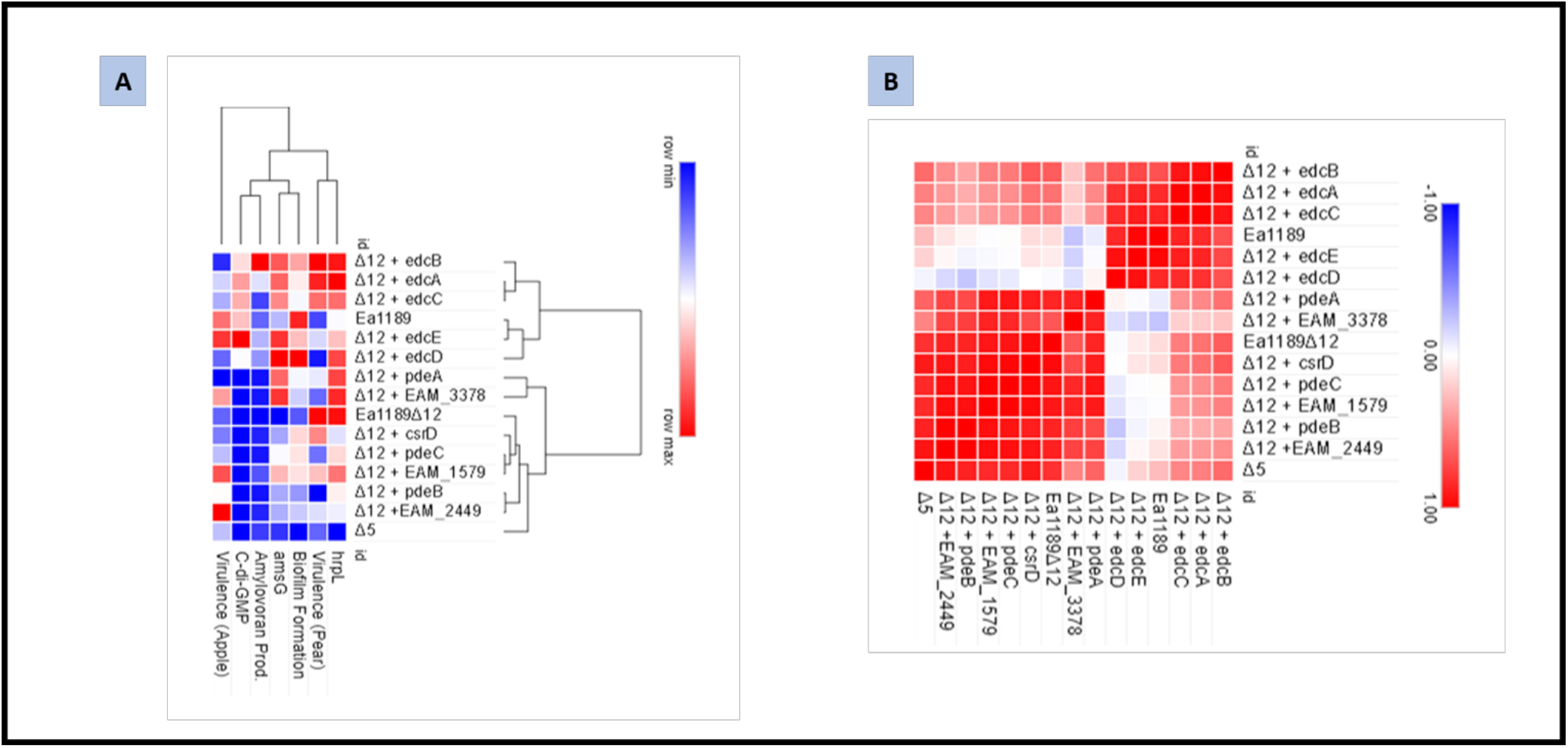
A) Aggregated data presenting several virulence factors including biofilm formation, amylovoran production, intracellular levels of c-di-GMP, virulence in apple shoots and immature pear fruit as wells as relative *hrpL* and *amsG* transcript levels (Relative to Ea1189) for *E. amylovora* Ea1189, Ea1189Δ5, Ea1189Δ12 and Ea1189Δ12 complemented with each individual deleted gene. The data presented in the form of a heatmap (using Morpheus) has been hierarchically clustered (Pearson correlation) (48) based on both the strain specificity and the evaluated factors. B) A similarity matrix computed based on the datapoints presented in panel A using Pearson correlation.

## Discussion

In this study, we used stepwise gene deletion to eliminate a c-di-GMP network from a pathogen in which this system has evolved to regulate transitions between T3SS-mediated and biofilm-mediated pathogenesis. Primarily in our study, we utilized Ea1189Δ12 as a representative model to study the effects of a systemic deletion of c-di-GMP metabolic components. Our aim was to evaluate the c-di-GMP metabolism-dependent and independent functions of each of the genetic components. However, owing to the possibility that the major driving force behind the observed phenotypic and transcriptomic changes in Ea1189Δ12 was the lack of intracellular production of c-di-GMP, we compared the phenotypic and transcriptomic profiles of Ea1189Δ5 (lacking the ability to synthesize c-di-GMP) against Ea1189Δ12 (lacking all proteins with GGDEF, EAL, or degenerate GGDEF and/or EAL domains). While both strains did not synthesize any c-di-GMP, their transcriptomes revealed several regulatory changes in fundamental processes including metabolism, substrate biosynthesis and ABC transport. This is indicative of larger systemic changes resulting from the deletion of the additional 7 genetic components of the c-di-GMP network that do not directly participate in c-di-GMP synthesis. Additionally, it is also noteworthy that the incremental changes occurring due to the shift from Ea1189Δ5 to Ea1189Δ12, do so in the absence of any detectable intracellular c-di-GMP. Phenotypically, Ea1189Δ5 contrasted Ea1189Δ12 in terms of reduced *hrpL* expression and virulence in the pear model. Reduced intracellular levels of c-di-GMP in *E. amylovora* were found to have an incremental effect on *hrpL* expression in vitro and increased virulence in planta (8). Combining these data, we were able to establish some baseline regulatory distinctions present in Ea1189Δ12 and Ea1189Δ5, and thus, corroborate the use of Ea1189Δ12 as a representative c-di-GMP network deletion strain.

Ea1189Δ12 displayed a loss in its ability to attach to a surface, even when allowed an extended contact time. Interestingly however, Ea1189Δ12 cells showed a high level of interaction with the surface, in terms of approaching the surface briefly, before retracting. Our results suggest that the flagellar filament (FliC) is responsible for the mitigation of this initial approach towards the surface and making initial contact. The overexpression of *hofC* (type IV pilus biogenesis assembly protein) enabled surface attachment in Ea1189Δ12, however this occurrence was contingent on the presence of FliC. While both the flagellum and the type IV pilus have been implicated in surface sensing and initial attachment, a critical limitation is the ability to sequentially categorize this occurrence. Our experimental setup evaluating attachment in real time collected images approximately every 5 to 8 seconds. This suggests that while both flagella and type IV pili might occur on the cell surface simultaneously, there is a functional distinction in their role during these initial stages. While our use of Ea1189Δ12 eliminated both the production of c-di-GMP and the loss of other related signaling components, the phenomena of surface sensing and attachment were found to be dependent on the presence of intracellular c-di-GMP, since the restoration of *edcA-E* led to the occurrence of surface attachment in Ea1189Δ12. In *P. aeruginosa*, c-di-GMP regulates type IV pilin biogenesis (23), and c-di-GMP production has been found to be further activated upon initial surface anchoring via the flagellum, leading to irreversible surface attachment and the initiation of multilayered biofilm formation (24). In *E. amylovora*, there appears to be two levels of regulation by c-di-GMP during initial surface contact. The existing presence of intracellular c-di-GMP is required for successful type IV pilus biogenesis, which although dispensable for initial surface contact will be required for surface anchoring.

In plant pathogenic bacteria, c-di-GMP regulates numerous pathways that contribute to virulence and host colonization (25). While biofilm formation has been shown to be positively affected by increasing intracellular levels of c-di-GMP in vitro in several systems including *E. amylovora* (8, 9), *Pseudomonas fluorescens* (26), *Xanthomonas oryzae* (27), and *Xylella fastidiosa* (28), a direct link between c-di-GMP mediated regulation and xylem colonization has not been established in these systems. An important finding in our study is that the absence of both intracellular c-di-GMP along with the elimination of the regulatory network involving c-di-GMP renders *E. amylovora* unable to colonize the xylem within apple shoots. In pear fruitlets, the T3SS is primarily utilized to infect host cells resulting in tissue necrosis, and the dependence on biofilm formation for successful pathogenesis is absent (29). Owing to the absence of c-di-GMP, *hrpL* transcription in vitro and virulence in immature pear fruit are elevated in Ea1189Δ12 compared to WT Ea1189. However, the infection of apple shoots involves the deployment of T3SS effectors (most importantly, DspE) within the apoplast to infect plant tissues, followed by attachment and biofilm formation within the xylem (15, 30). The translocation of DspE is negatively regulated by c-di-GMP in *E. amylovora* (9). Despite this, the apoplastic colonization by Ea1189Δ12 within apple leaf tissue appeared to be relatively weak. In terms of biofilm formation within the xylem, evidence from our previous study suggests that a reduction in intracellular c-di-GMP levels due to the simultaneous deletion of multiple *edc* genes resulted in increased tissue necrosis inclusive of the petiole in *E. amylovora* (8). This suggests that the reliance on c-di-GMP for xylem colonization is dependent on both the generation of intracellular c-di-GMP as well as the complex regulation of physical biofilm formation through multigene components in *E. amylovora*.

The process of biofilm formation is composed of distinct stages including surface sensing, initial surface attachment followed by irreversible surface attachment, the multiplication of cells to form multilayered biofilms and eventually dispersal of cells from the biofilm (31). Our experimental setup to evaluate the transcriptomic profiles of *E. amylovora* during these various stages of biofilm formation aimed at deciphering the regulatory pathways affected by c-di-GMP in each of these stages (barring the dispersal stage). However, due to the inability of Ea1189Δ12 to attach to a surface or develop biofilms, our sampling for this strain was limited to the cells that were surface exposed but unattached (Unbound sample type). With this consideration, our comparison of DEGs overall revealed that the greatest level of transcriptomic heterogeneity between Ea1189Δ12 and Ea1189 occurred in the Unbound sample type, and this level of heterogeneity decreased to the lowest level within the Ea1189_Bound vs. Ea1189_Biofilm comparative metric. A majority of the DEGs were a part of metabolic pathways, both in the Unbound comparative metric as well as the other conditions. While studies focused on the transcriptomic profiling of biofilm formation are abundant among bacterial pathogens, the bifurcation of biofilm stages in this process is less prevalent. Kang et al. (32) used laser microdissection to segregate portions of a *P. aeruginosa* biofilm based on their location relative to the surface or the liquid-air interface. Their results suggested that the transcriptomic profile in each section of a biofilm was significantly different, with varied DEGs present in each sectional sample (32). The RNA-Seq profiling of *X. fastidiosa* cells grown in microfluidic chambers to study the effect of calcium on cellular function revealed that the largest characterized subgroup of DEGs contributed to metabolic functions (33). Overall, our results indicate that *E. amylovora* has evolved to rely on c-di-GMP based transcriptional regulation to shift its metabolic profile to transition into biofilm formation, and this regulation begins to occur as soon as cells come in contact with a surface and will adjust during the course of biofilm initiation, tapering off when cells are in a mature biofilm.

Our aggregate data analysis of multiple virulence factors in *E. amylovora* Ea1189, Ea1189Δ12, and single gene complement strains, revealed distinct patterns of c-di-GMP dependent and independent regulation based on the presence or absence of c-di-GMP in the strains. Biofilm formation, *amsG* expression, and amylovoran production were primarily elevated in strains that were able to produce c-di-GMP. These results are consistent with the results observed in previous studies when the deletion of *pdeA-C* resulted in elevated intracellular levels of c-di-GMP, resulting in increased EPS production and biofilm formation (9).. So far, no c-di-GMP dependent receptors that regulate amylovoran production have been identified in *E. amylovora*. Virulence regulation overall showed a more diffuse pattern in terms of the impact of the complemented genes. For each of the three analyzed criteria including *hrpL* expression, and necrosis in immature pear and apple shoots, c-di-GMP levels were not the sole limiting factor. Ea1189Δ12 displayed elevated virulence in pear tissue, however, the inability to form biofilms within the xylem limited its virulence in apple shoots. Thus, counterintuitively, although *hrpL* transcription is reduced with the addition of c-di-GMP, the same factor results in an increased ability to form biofilms, thus translating into elevated virulence in apple shoots. Of all the Edcs, EdcE was found to most strongly downregulate both *hrpL* expression and virulence in pears. In apple shoots, EAM_2449, EdcE, and CsrD led to the greatest increase in necrosis (primarily indicative of xylem colonization). Although EAM_2449 and CsrD don’t enzymatically contribute to the generation of c-di-GMP, the presence of GGDEF and/or EAL motifs in these proteins could be a target for c-di-GMP binding, leading to feedback regulation (34, 35). While the mechanism of how *hrpL* is transcriptionally regulated by c-di-GMP is not understood in *E. amylovora*, in the related phytopathogen *Dickeya dadantii*, T3SS regulation is regulated by the sRNA chaperone Hfq through the repression of two Dgcs, GcpA and GcpL (36). In addition, the flagellar master regulator FlhDC also regulates the T3SS by exerting negative pressure on c-di-GMP levels through the activation of the Pde EcpC (37). In *E. amylovora*, although Hfq has been linked with T3SS regulation, the link to the intermediary being c-di-GMP is absent.

Overall, our study highlights the evolutionary importance of c-di-GMP in the *E. amylovora* disease cycle. Our approach of eliminating all Dgc and Pde components in *E. amylovora* and systematically restoring them highlighted the dual modality of control imposed by this overall system, including both c-di-GMP dependent and independent regulation of virulence orchestration in the host. Interaction with the host, specifically in terms of xylem colonization was found to be a complex process that necessitates a metabolic shift in cells, a process heavily dependent on c-di-GMP. The study highlighted several new genes that strongly regulate amylovoran production and T3SS. Dissecting the specific pathways by which these genes impact virulence will be critical to further expanding the known c-di-GMP regulatory network in *E. amylovora*.

## Materials and Methods

### Bacterial strains, plasmids and growth conditions

All bacterial strains, and plasmids used in this study along with their relevant characteristics are described in Table S1. Unless specified otherwise, *E. amylovora* strains were grown in Luria-Bertani (LB) based medium amended with one or more of the appropriate antibiotics: ampicillin (Ap; 100 µg/ml), chloramphenicol (Cm; 10 µg/ml), gentamicin (Gm; 10 µg/ml) or kanamycin (Km; 100 µg/ml). Overexpression vectors (*hofC* OE and *fliC* OE) were induced using 1mM isopropyl-b-D-thiogalactopyranoside (IPTG).

### Genetic manipulation and bioinformatics

Reference genome sequence for *E. amylovora* (Accession: ATCC-49946/FN666575) was obtained from NCBI. Artemis (Java) was used to browse the genome. The standard lambda Red recombinase protocol was used to construct chromosomal deletion mutants(38). To complement the individual deleted genes, the full length sequences of the genes were briefly amplified, and the purified gene fragments were transformed into Ea1189Δ12 harboring pKD46 (induced with arabinose) along with a retained Cm^R^/Km^R^ cassettes (originally amplified from pKD3/pKD4 source plasmids) in the target gene being restored. Transformed cells were recovered after an 18h incubation and were screened for a loss of the resistance cassette, followed by sanger sequencing to confirm the replacement of the originally deleted gene.

### Confocal microscopy to monitor attachment and biofilm formation in flow cells

To monitor initial surface interaction and attachment, *E. amylovora* strains expressing pMP2444::*gfp* (16) were grown for 18 h at 28°C and normalized to an OD_600_ of 0.5. A total of 1 ml of inoculum for each strain was introduced into a flow cell chamber in a µ-Slide VI 0.5 glass bottom slide (Ibidi, Martinsried, Germany). Immediately, the base of the flow chamber was repeatedly imaged using a Olympus FlouView 1000 confocal laser scanning microscope (Olympus, MA, USA). Images were acquired for up to 1 h or until the frame was saturated with fluorescent cell signals. Following this, the flow cell chamber was flushed with 5 ml of 0.5X phosphate buffered saline (PBS). To evaluate biofilm formation, following the initial attachment incubation, the flow chamber was subjected to flow (0.5X LB) using a peristaltic pump (Ismatec REGLO Digital 4-CH pump) (Cole-parmer IL, USA) for 5 h. Fluorescent Z-stacked images were acquired to measure overall attachment and biofilm levels in the flow cell chambers (16). ImageJ software was used to process these images and to graph the GFP signal intensity profile for the Z-stacked images (39).

### Scanning electron microscopy to monitor xylem colonization in apple shoots

Apple shoots were inoculated as previously described (8). Strains were grown overnight at 28°C and normalized to an OD_600_ of 0.2. Scissors dipped in inoculum were used to make an incision between two peripheral veins on young apple leaves (*Malus x domestica* cv. Gala on M9 rootstock). At 5 dpi, inoculated leaves along with the attached petiole were harvested. Cross sections of the petioles and apoplast tissue were fixed using 2.5% paraformaldehyde-2.5% glutaraldehyde, followed by ethanol dehydration at increasing concentrations as previously described. Samples were imaged using the JEOL 6610LV (Japan Electron Optics Laboratory Ltd., Tokyo, Japan).

### RNA-Seq sample acquisition, sequencing, and data analysis

WT *E. amylovora* Ea1189, Ea1189Δ5 and Ea1189Δ12 were grown overnight in LB medium, sub-cultured and harvested at the mid-log phase prior to RNA extraction. To collect cell samples from various stages of biofilm development, the protocol described in this study to monitor biofilm formation in flow cells remained largely the same (note that the strains were not fluorescently labelled). Inoculum injected into the flow chamber for the strains was collected 1 h after the cells were allowed to interact with the surfaces in the flow chamber (Unbound sample). Following this, the flow chamber was flushed with PBS as described and the cells that remained attached to the surfaces in the flow chamber were collected to represent the ‘Bound sample’. Flow driven biofilm formation was allowed to proceed for 24 h before collecting attached cells representing ‘Biofilm sample’. For RNA extraction, as per a previously described protocol (40), cells were washed with 0.1% N-lauryl sarcosine sodium salt, followed by treatment with the lysis buffer (1% SDS in 10 mM EDTA and 50 mM sodium acetate, pH 5.1) and a 5 min incubation in boiling water. The extracted RNA was treated for residual DNA contamination using the TURBO DNA-free kit (Thermo Fisher Scientific, MA, USA), and concentrated using the RNA Clean and Concentrator-25 kit (Zymo Research, CA, USA). The ‘Unbound’ samples were treated after their collection from the flow chamber, for the other two sample types, the initial lysis and wash steps were conducted by injecting the buffers directly into the flow chamber and suctioning the fluid out.

RNA samples were analyzed for quality control on the Agilent 4200 TapeStation (Agilent Technologies, CA, USA). The QIAseq FastSelect 5S/16S/23S rRNA removal kit was used to treat samples prior to library prep with the TruSeq Stranded Total RNA Library Prep Kit (Illumina, CA, USA). Sequencing was conducted on the Illumina HiSeq 4000 at 50 bp single-end reads.

For data analysis, adaptor barcodes were filtered using Trimmomatic v 0.36 (single end criteria: ILLUMINACLIP:TruSeq3-SE:2:30:10 LEADING:3 TRAILING:3 SLIDINGWINDOW:4:15) (41). Trimmed sequences were mapped to the *E. amylovora* ATCC-49946 genome using Bowtie v 2.4.1. HTSeq v 0.11.2 (42, 43). Differential expression analysis was conducted using DESeq2 v 3.12 with an FDR cutoff of 0.05 and a minimum accepted fold change of 2 (44). Gene Ontology enrichment analysis visualization was conducted using iDEP 9.1 tool (45).

### Measuring biofilm formation

Biofilm formation in vitro was quantified using a previously described protocol (46). Strains were grown overnight at 28°C, normalized to an effective OD_600_ of 0.05 in 0.5X LB. A polypropylene bead (diameter 7mm) (Cospheric LLC, CA, USA) was suspended in this inoculum and incubated for 48 h at 28°C. The beads were then stained with a 0.1% crystal violet solution, which was solubilized using a combination of 40% methanol and 10% glacial acetic acid. Data was collected in the form of the OD_595_ of the elution. Data includes three biological replicates, all processed using JMP statistical software™; this software was used for all phenotypic numerical data analysis.

### Measuring amylovoran production

Amylovoran production was quantified using a turbidometric assay as previously described(9). Strains were grown overnight in LB and their OD_600_ was normalized. Strains were then grown in MBMA medium (with sorbitol) for 48 h at 28°C. Cetylpyridinium chloride (CPC at 50mg/ml) was used to precipitate amylovoran from the supernatant of the incubated cultures. Data is presented as CPC precipitation levels normalized by cell density.

### q-RT-PCR to measure gene expression

To measure *amsG* and *hrpL* expression, strains were grown overnight in LB at 28°C. Cell cultures were then washed and resuspended in MBMA and HRP-MM medium respectively and incubated for 6h. To validate RNA-Seq data, identical sample treatment and collection protocols were used, with the exception of measuring transcript levels only for representative gene targets. RNA extraction and concentration for all samples were conducted using the protocols described for RNA-Seq sample collection. cDNA was synthesized using the High Capacity RT kit (Applied Biosystems, CA, USA). SYBR green PCR master mix (Applied Biosystems, CA, USA) was used for quantitative PCR experiments. *recA* was used as an endogenous control. The delta C_T_ method was used to compare transcriptomic fold changes (47). Three biological replicates were included in the studies.

### Quantifying intracellular levels of c-di-GMP

Intracellular levels of c-di-GMP were quantified as previously described(8). Strains were grown in overnight in LB, sub-cultured, collected at mid-log phase and lysed (with 40% acetonitrile and 40% methanol) at -20°C for 1 h. Relative levels of c-di-GMP in the samples was established against a standard curve generated using synthesized c-di-GMP (Axxora Life Sciences Inc., CA, USA) using a Quattro Premier XE instrument (Waters Corp. MA, USA).

### Virulence assays

Relative virulence levels of strains were compared using immature pear fruits as previously described (8). Immature pears were stab inoculated with cultures at a concentration of 10^4^CFU/ml and incubated for 5 days at 28°C under high humidity. Data was collected in the form of the diameter of necrotic lesions formed on the pears. Apple shoots were inoculated using the same described protocol used to evaluate xylem colonization through SEM. Data was collected in the form of necrotic lesion length along the shoot 8 dpi.

## Acknowledgements

We would like to acknowledge the assistance provided by the MSU RTSF core in RNA-Seq sample prep, along with Carol Flegler and Melinda Frame at the MSU Microscopy core for their assistance with SEM and CLSM.

## Notes

### Competing Interest Statement

The authors have declared no competing interest.

