## Supplemental Figures and Table for "The cyclic di-GMP network is a global regulator of phase-transition and attachment-dependent host colonization in *Erwinia amylovora*"

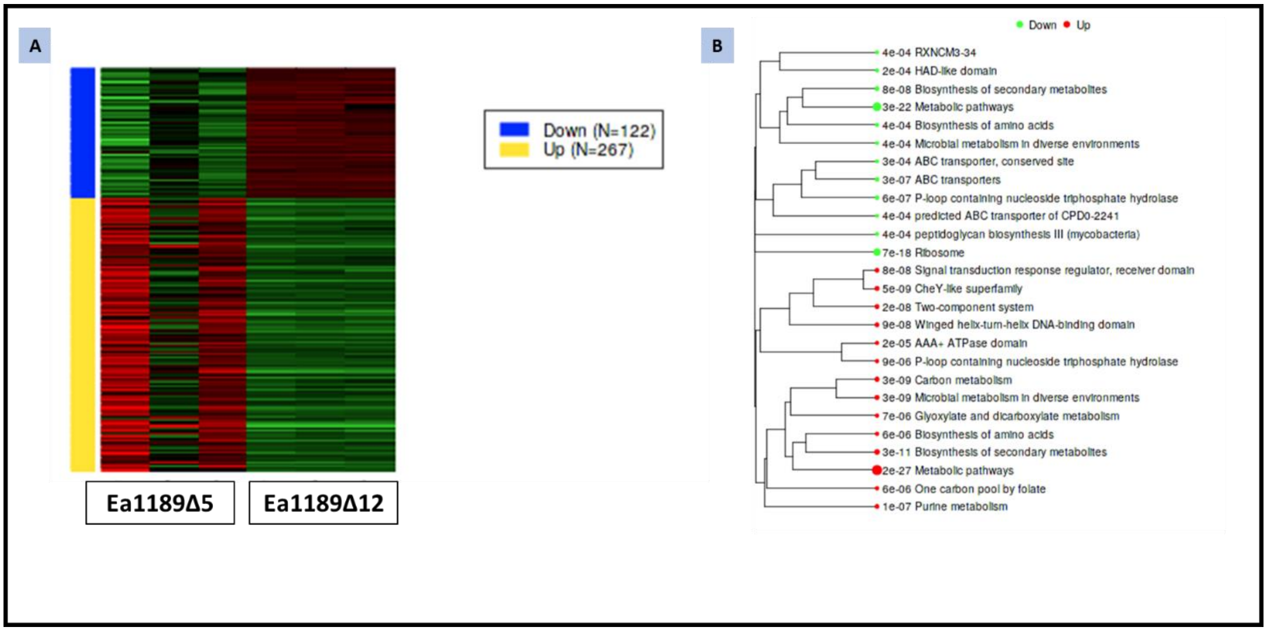


Supplementary Figure 1: A) A clustered representation of the up and down regulated DEGs in Ea1189Δ12 compared to Ea1189Δ5. B) A hierarchical clustering of the GO based enriched regulatory pathways represented by the DEGs for Ea1189Δ12 vs Ea1189Δ5. The intensity of the branch markings signifies the size of the gene set contained within each identifier. iDEP was used for data visualization and GO enrichment analysis (45).

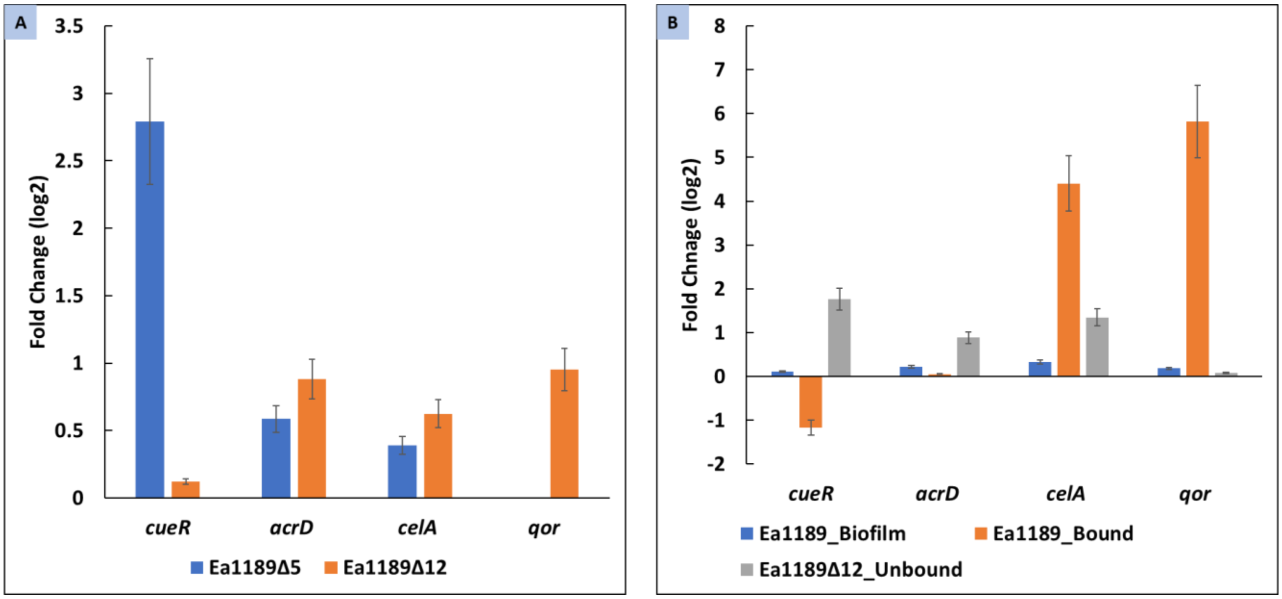


Supplementary Figure 2: q-RT-PCR was used to validate the RNA-Seq results using representative genes. The delta $C_T$ method was used to calculate the relative fold change in expression levels for the gene targets relative to Ea1189 (A) andEa1189_Unbound condition (B)(47). Error bars indicate standard error of the means.

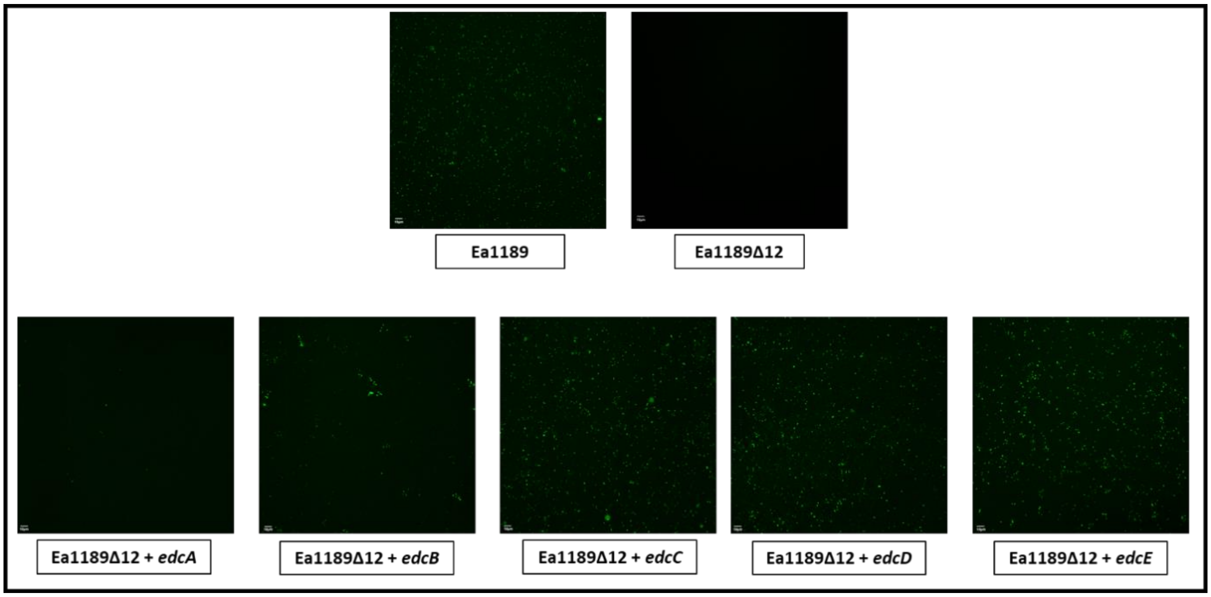


Supplementary Figure 3: Z-stacked confocal scanning microscopy images displaying the level of biofilm formation under microfluidic flow conditions for Ea1189, Ea1189Δ12 and Ea1189Δ12 complemented with genes *edcA-E*. Relative to Ea1189Δ12 is severely impaired in its ability to form biofilms within the flow chamber. The restoration of *edcA-E* genes results in an elevation of intracellular c-di-GMP levels, thus enabling Ea1189Δ12 to attach and form biofilms under flow.

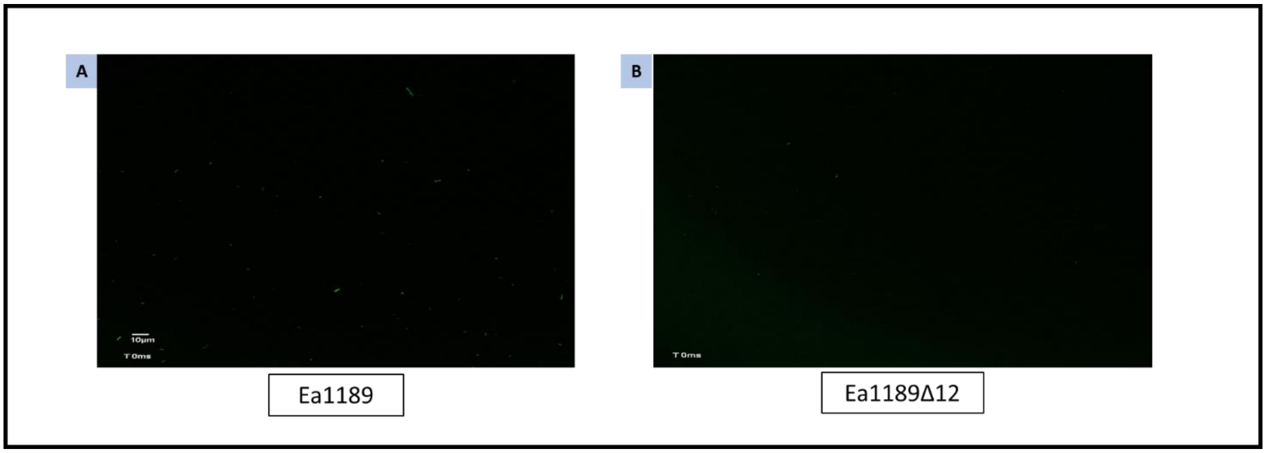


**Please note that the videos for supplementary figure 4A and 4B have been uploaded as separate files to the submission server.**

Supplementary Figure 4: Videos documenting the initial contact of Ea1189 (A) and Ea1189Δ12 (B) cells with the inner surface of a flow cell chamber upon injection through the flow cell inlet. Confocal microscopy images of GFP labelled cells of both strains were captured at the base of the flow cell chamber at the same fixed location approximately in the middle of the chamber. Compiled images for each variant are presented as a time lapse video. Upon approaching the basal surface of the flow chamber, Ea1189 cells were able to attach to the surface and retain attachment over time, leading to a visual saturation of fluorescent cells within the frame of the image. However, Ea1189Δ12 cells were found to approach the basal surface (as evidenced by the flickering GFP signals within the video), however, these cells failed to attach to the surface over time.

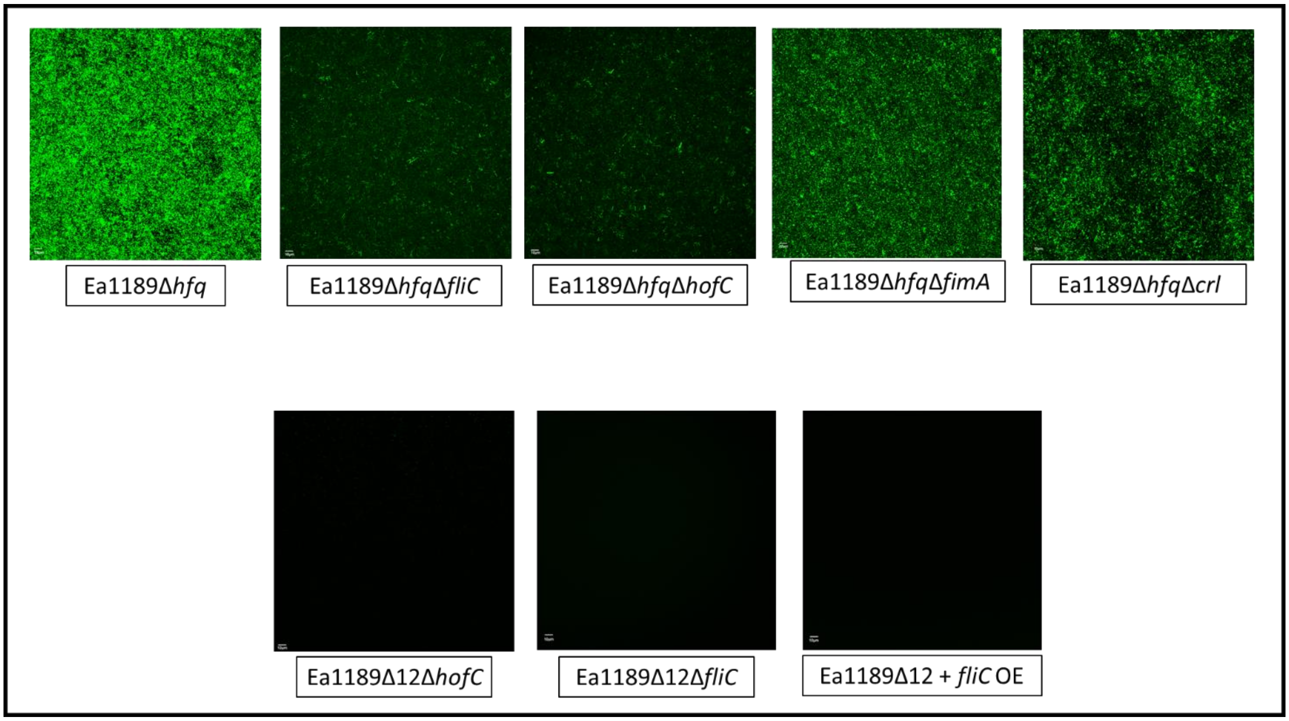


Supplementary Figure 5: Confocal Z-stack images depicting the level of attachment within a flow chamber after a 1h incubation period for Ea1189Δ*hfq* and double mutants targeting the flagellum (*fliC*), type IV pilus (*hofC*), fimbriae (*fimA*) and curli fimbriae (*crl*). The deletion of *fliC* and *hofC* in Ea1189Δhfq had the highest degree of negative impact on attachment within the flow chamber. The overexpression of *fliC* in Ea1189Δ12 did not yield any attachment. As negative controls, we included Ea1189Δ12Δ*hofC* and Ea1189Δ12Δ*fliC* in this sample set.

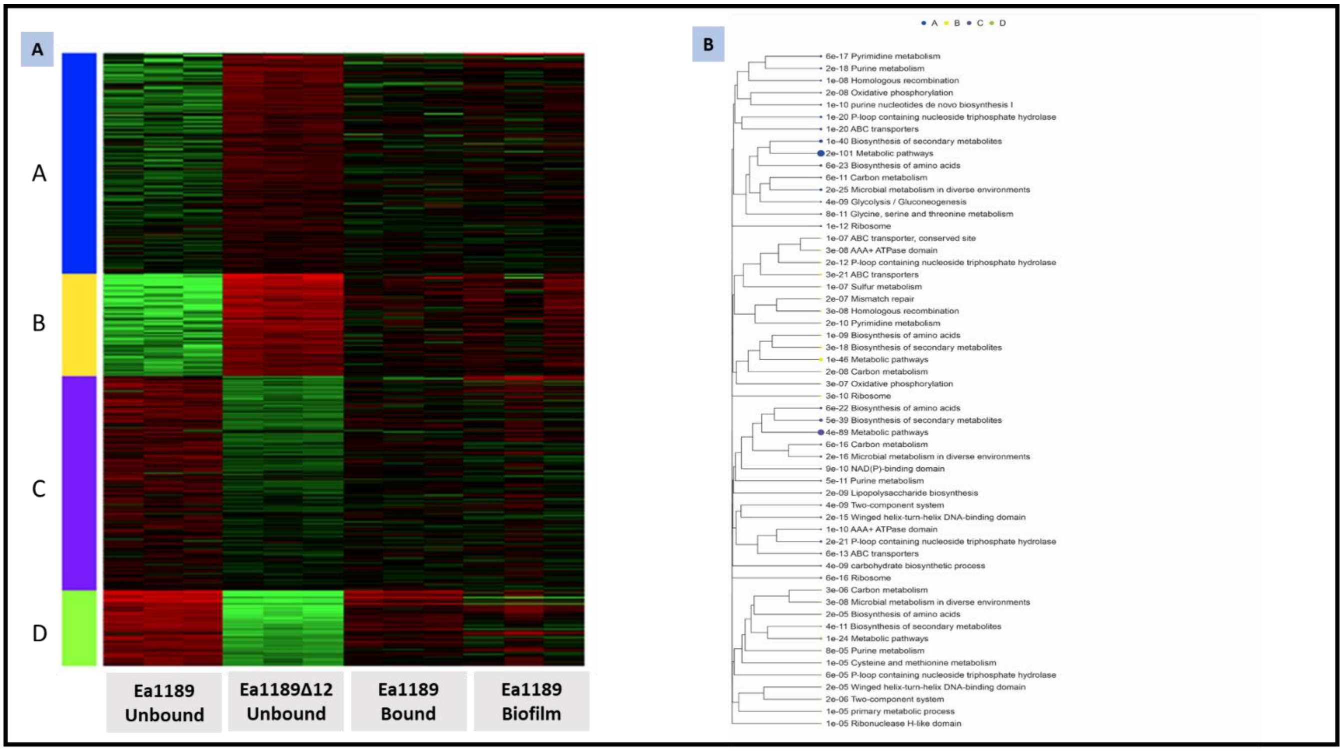


Supplementary Figure 6: A) A clustered heatmap for Ea1189 and Ea1189Δ12 compared during various stages of biofilm development by RNA-Seq. The DEGs are clustered into four sub-groups by treatment sample type. The letters to the left of the heatmap are arbitrary cluster IDs. The gene ontology distribution tree highlights the cellular pathways in which the DEGs for each of the clusters are enriched in. The color coding is indicative of the cluster ID and the size of the circular dot is a relative comparison of the numerical abundance of DEGs in a specific pathway. iDEP software was used to conduct this visual analysis (44).

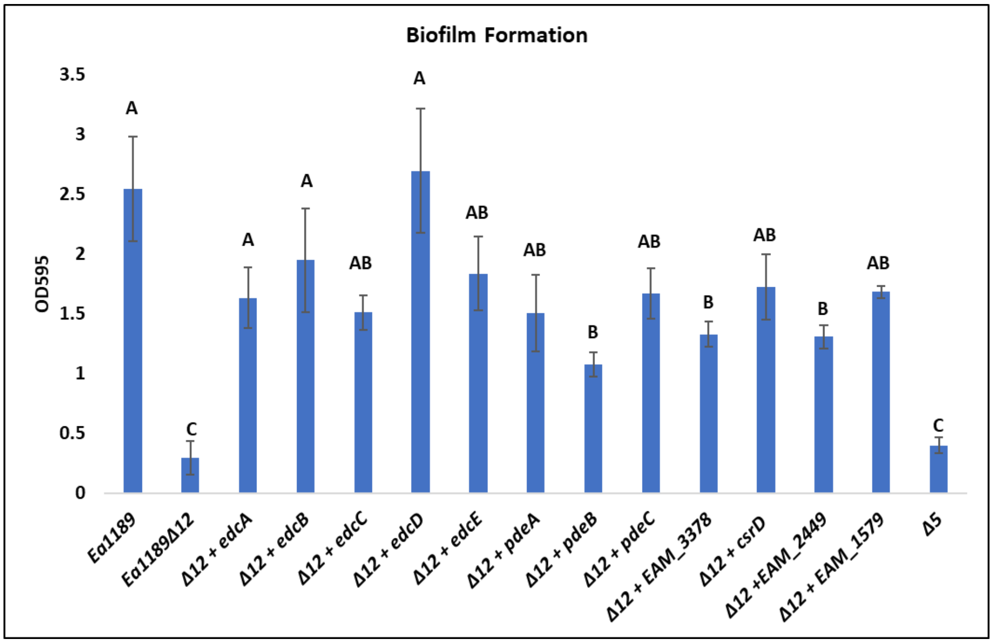


Supplementary Figure 7: Relative levels of biofilm formation for Ea1189, Ea1189Δ5, Ea1189Δ12 and Ea1189Δ12 complemented with each individual deleted genetic component. The data is represented in the form of the $OD_{595}$ for crystal violet binding. The error bars represent standard errors of the means. The significance letters above the bars are based on statistically significant differences ($P<0.05$) calculated by Tukey’s HSD.

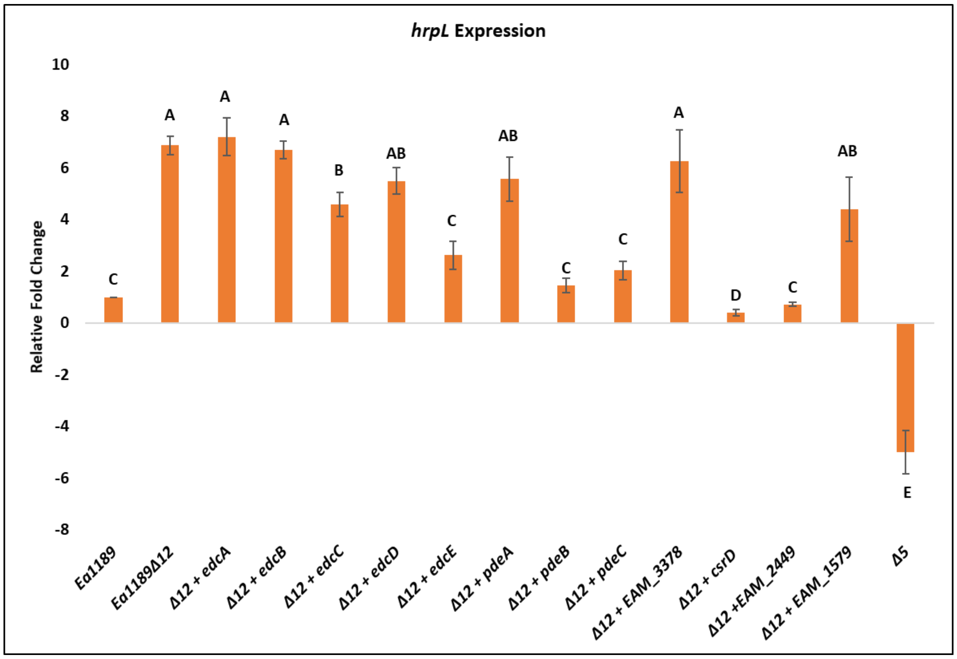


Supplementary Figure 8: Relative gene expression levels for *hrpL* compared for Ea1189Δ5, Ea1189Δ12 and Ea1189Δ12 complemented with each individual deleted genetic component against WT Ea1189. The delta $C_T$ method was used to process the data (47). The error bars represent standard errors of the means. The significance letters above the bars are based on statistically significant differences (P<0.05) calculated by Tukey’s HSD.

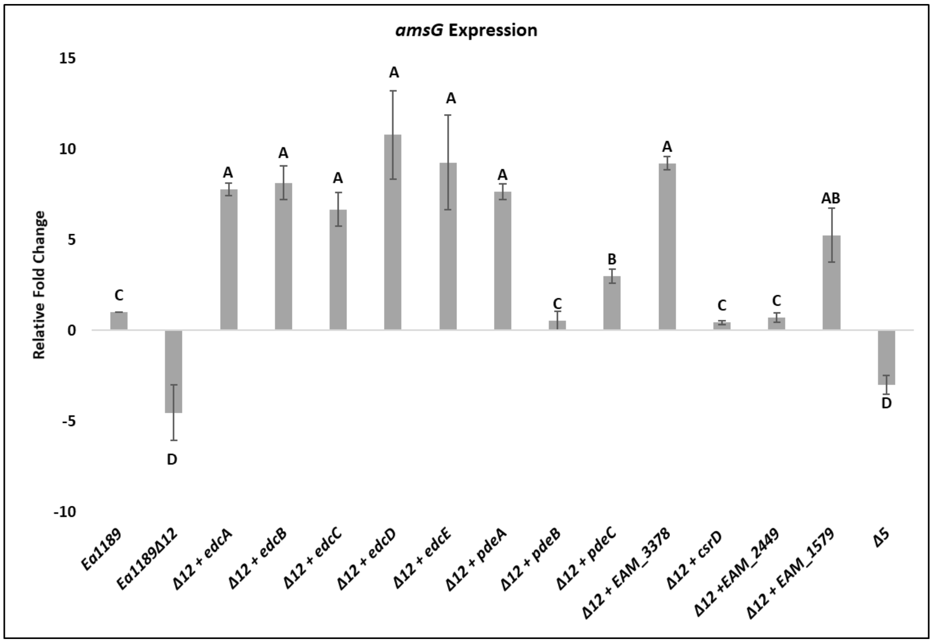


Supplementary Figure 9: Relative gene expression levels for *amsG* compared for Ea1189Δ5, Ea1189Δ12 and Ea1189Δ12 complemented with each individual deleted genetic component against WT Ea1189. The delta $C_T$ method was used to process the data (47). The error bars represent standard errors of the means. The significance letters above the bars are based on statistically significant differences ($P<0.05$) calculated by Tukey’s HSD.

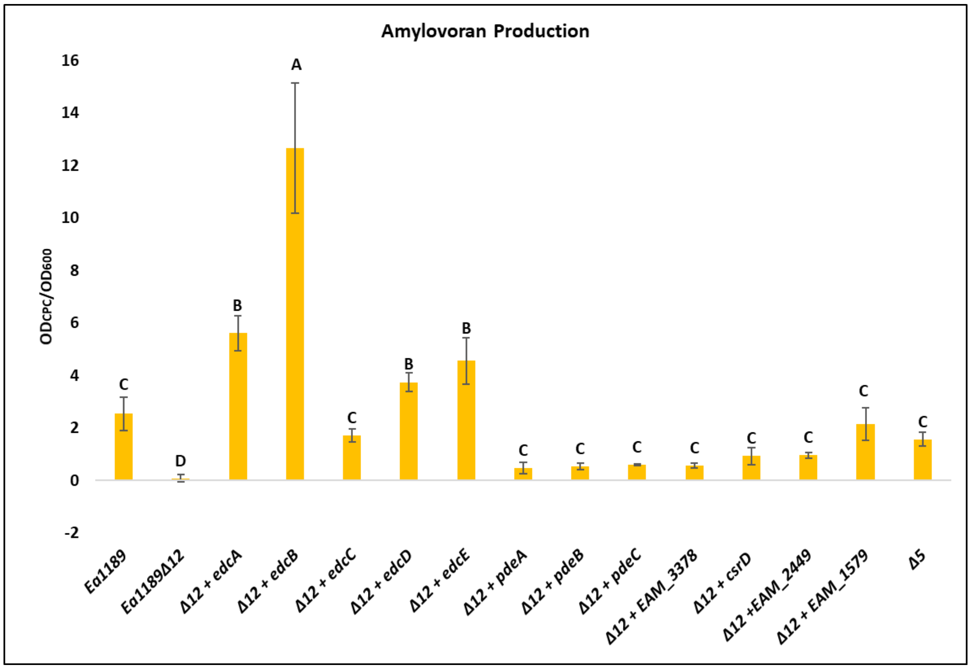


Supplementary Figure 10: Relative levels of amylovoran production for Ea1189, Ea1189Δ5, Ea1189Δ12 and Ea1189Δ12 complemented with each individual deleted genetic component. The data is represented in the form of the $OD_{600}$ for CPC binding, normalized by cell density. The error bars represent standard errors of the means. The significance letters above the bars are based on statistically significant differences ($P<0.05$) calculated by Tukey's HSD.

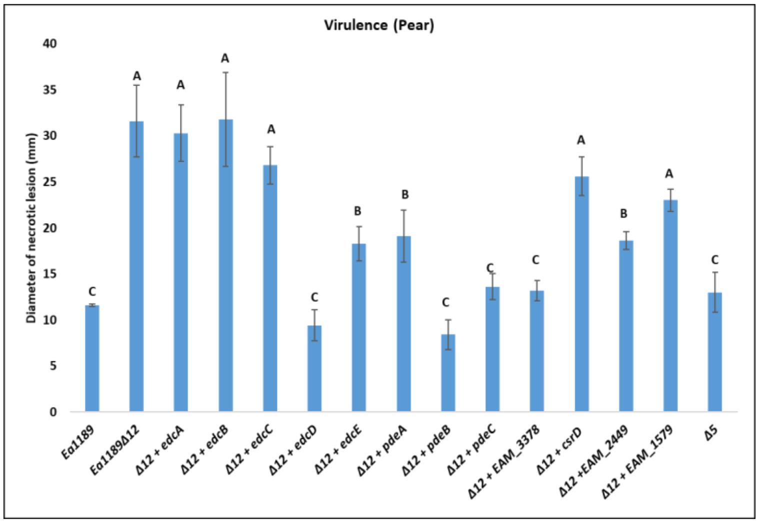


Supplementary Figure 11: Virulence levels assayed in the immature pear model for Ea1189, Ea1189Δ5, Ea1189Δ12 and Ea1189Δ12 complemented with each individual deleted genetic component. The data is represented in the form of measurable necrotic lesion diameters on the infected pear surface. The error bars represent standard errors of the means. The significance letters above the bars are based on statistically significant differences ($P<0.05$) calculated by Tukey’s HSD.

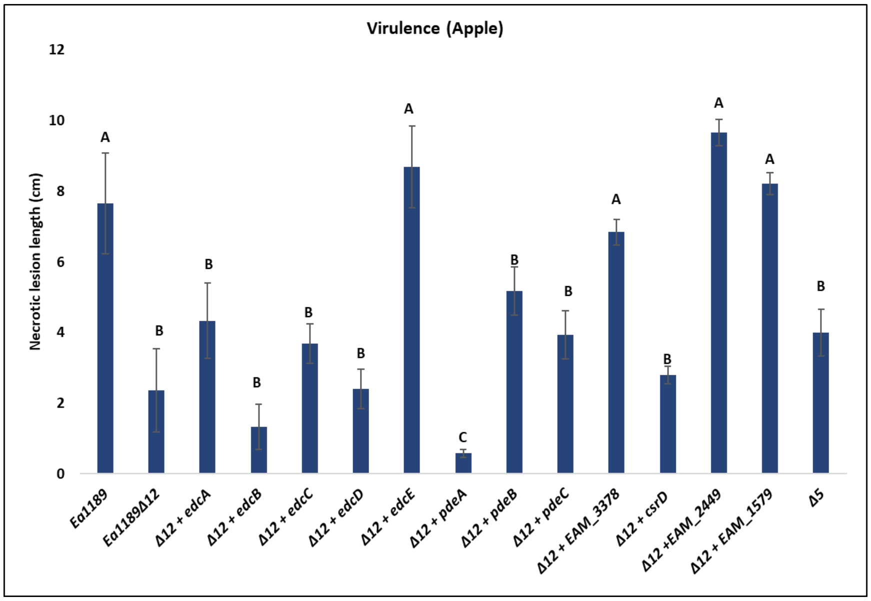


Supplementary Figure 12: Virulence levels assayed in the apple shoot model for Ea1189, Ea1189Δ5, Ea1189Δ12 and Ea1189Δ12 complemented with each individual deleted genetic component. The data is represented in the form of measurable necrotic lesion lengths along the leaf and shoot during infection. The error bars represent standard errors of the means. The significance letters above the bars are based on statistically significant differences (P<0.05) calculated by Tukey's HSD.

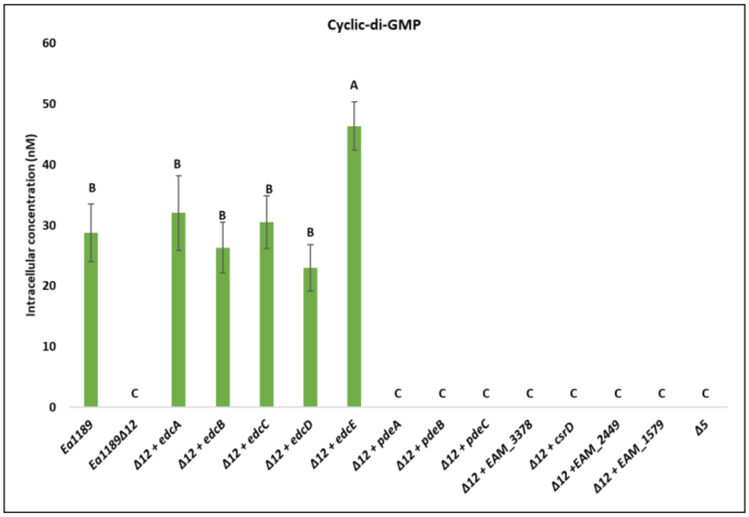


Supplementary Figure 13: Intracellular levels of c-di-GMP for Ea1189, Ea1189Δ5, Ea1189Δ12 and Ea1189Δ12 complemented with each individual deleted genetic component as measured by LC-MS-MS. The error bars represent standard errors of the means. The significance letters above the bars are based on statistically significant differences (P<0.05) calculated by Tukey’s HSD.

Supplementary Table 1

| Strain/Plasmid | Relevant Characteristics | Source |
|---|---|---|
| *E. amylovora* strains | | |
| Ea1189 | Wild Type | (8) |
| Ea1189Δ5 | Deletion of *edcA-E* | Sundin lab collection |
| Ea1189Δ12 | Deletion of *edcA-E*, *pdeA-C*, EAM_3378, EAM_3136, EAM_2449 and EAM_1579 | This study |
| Ea1189Δ12Δ*fliC* | Deletion of *fliC* in Ea1189Δ12 | This study |
| Ea1189Δ12Δ*hofC* | Deletion of *hofC* in Ea1189Δ12 | This study |
| Ea1189Δ12 + *edcA-E/pdeA-C/EAM_3378/csrD/EAM_2449/EAM/1579* | Chromosomal restoration of the indicated gene in Ea1189Δ12 | This study |
| Ea1189Δ*hfq* | *hfq* deletion mutant | (21) |
| Ea1189Δ*hfq*Δ*fliC/hofC/fimA/crl* | Double deletion mutant of *hfq* and the indicated gene | This study |

| Plasmids | | |
|---|---|---|
| pKD3 | $Cm^r$ cassette flanking FRT* sites; $Cm^r$ | (38) |
| pKD4 | $Km^r$ cassette flanking FRT sites; $Km^r$ | (38) |
| pKD46 | L-Arabinose-inducible lambda red recombinase; $Ap^r$ | (38) |
| pTL18 | IPTG-Inducible FLPase, $Tet^R$ | (49) |
| pBBR1-MCS5 | Broad-host-range cloning vector**; R6K ori; $Gm^r$ | (50) |
| pMP2444 | pBBR1MCS-5 expression *gfp* under *lac* promoter, $Gm^r$ | (51) |
| pEVS143 | Broad-host-range, IPTG inducible (Ptac) cloning vector; inducible $Cm^r$ and GFP $Km^r$ | (52) |
| *hofC* OE | *hofC* in pEVS143 | This study |

| | | |
|---|---|---|
| *fliC* OE | *fliC* in pEVS143 | This study |

*FRT: Flippase target recognition; **MCS: Multiple cloning site.

Supplemental Files: DEG list.csv: Differentially expressed gene list for all sample comparison metrics along with log2 fold change and adjusted P-values.

GO enrichment gene list.csv: Gene ontology pathway enrichment and included genes in the overall clustered analysis of DEGs.
